# Sexual selection moderates heat stress response in males and females

**DOI:** 10.1101/2022.03.20.485015

**Authors:** Maria Moiron, Lennart Winkler, Oliver Yves Martin, Tim Janicke

**Affiliations:** Centre d’Écologie Fonctionnelle et Évolutive, CNRS, Univ Montpellier, EPHE, IRD, 1919 Route de Mende, 34293 Montpellier Cedex 05, France; Institute of Avian Research, Wilhelmshaven, Germany; Applied Zoology, Technical University Dresden, Zellescher Weg 20b, 01062 Dresden, Germany; Department of Biology & Institute of Integrative Biology IBZ, ETH Zurich, Universitätstrasse 16, 8092 Zürich, Switzerland

**Keywords:** heatwaves, mating system, reproductive success, thermal stress, *Tribolium castaneum*

## Abstract

A widespread effect of climate change is the displacement of organisms from their thermal optima. The associated thermal stress imposed by climate change has been argued to have a particularly strong impact on male reproduction but evidence for this postulated sex-specific stress response is equivocal. One important factor that may explain intra- and interspecific variation in stress responses is sexual selection, which is predicted to magnify negative effects of stress. Nevertheless, empirical studies exploring the interplay of sexual selection and heat stress are still scarce. We tested experimentally for an interaction between sexual selection and thermal stress in the red flour beetle *Tribolium castaneum* by contrasting heat responses in male and female reproductive success between setups of enforced monogamy *versus* polygamy. We found that polygamy magnifies detrimental effects of heat stress in males but relaxes the observed negative effects in females. Our results suggest that sexual selection can reverse sex differences in thermal sensitivity, and may therefore alter sex-specific selection on alleles associated with heat tolerance. We argue that these findings have important implications for predicting the role of sexual selection for the adaptation to current global warming and increased frequency of extreme climatic events.

## Introduction

Ambient temperature is one of the most variable environmental factors that terrestrial organisms face during their lifetimes and is predicted to have profound effects on behaviour, physiology and life-history traits (Angilletta 2009; Huey *et al*. 2012; Tattersall *et al*. 2012). This is particularly true for ectotherms in which ambient temperature has a strong influence on body temperature and thus on the physiology of an individual (Huey & Kingsolver 1993). There is an increased awareness of intensifying global warming associated with an increasing frequency of extreme climatic events, which both are expected to further accelerate the ongoing extinction crisis (Bellard *et al*. 2012; Ummenhofer & Meehl 2017). Hence, the study of how organisms are affected by and adapt to changes in ambient temperature has become a central topic in ecology and evolutionary biology over the last two decades (Parmesan 2006; Walsh *et al*. 2019). The Intergovernmental Panel on Climate Change is currently predicting a rise of global surface temperature of 1.2 to 3.0 °C until 2060 (IPCC 2021), which will be likely linked to a higher frequency of extreme weather phenomena such as drought and heatwaves (Coumou & Rahmstorf 2012; Fischer & Knutti 2015). Despite the current interest in understanding the effects of climate change on the biosphere, our knowledge on how heat stress affects reproductive performance and the selective processes that modulate heat responses is still limited. As such, a comprehensive understanding of the different evolutionary processes driving adaptation to heat stress is presently of vital importance.

One prominent but still largely unexplored evolutionary process that may modulate heat responses is sexual selection (Garcia-Roa *et al*. 2020; Pilakouta & Ålund 2021), here defined as selection arising from competition for mating partners and/or their gametes (Jones & Ratterman 2009). On an evolutionary scale, theory predicts that sexual selection is a major driver of sex differences in morphology, physiology and life-history traits (Schärer, Rowe & Arnqvist 2012; De Lisle 2019). More specifically, sexual selection is often found to be stronger on males compared to females (Janicke *et al*. 2016), which may translate into stronger purifying selection against deleterious alleles on males (Winkler *et al*. 2021), and may give rise to sex-specific reproductive strategies (Schärer, Rowe & Arnqvist 2012). Importantly, sexual selection may select for a higher allocation of resources into current reproduction at the cost of future fitness. In particular, for species with stronger sexual selection on males, the so-called “live fast die young” strategy predicts that males invest less resources into traits facilitating survival including health and stress resilience (Bonduriansky *et al*. 2008; Tarka *et al*. 2018), in favour of an enhanced reproduction. Therefore, sexual selection is predicted to promote the evolution of higher susceptibility to environmental stress in males compared to females (Hämäläinen *et al*. 2018). However, formal theoretical work in support of this prediction is scarce and sexual selection on condition-dependent traits may mitigate the postulated sex differences as explored, for example, for sex-specific immunocompetence (Stoehr & Kokko 2006).

In addition to the predicted evolution of sex-specific life histories, sexual selection may amplify the negative effects of environmental stress, because intra-sexual competition for mating partners is predicted to favour individuals with superior condition, defined as the pool of acquired resources that can be allocated towards various fitness routes including traits conveying a higher competitiveness at pre- and postcopulatory episodes of sexual selection. The so-called “genetic capture” hypothesis posits that condition is a highly polygenic trait and that sexual selection on condition-dependent sexual traits favours genetic variants that are also favoured by natural selection in a given environmental context (Rowe & Houle 1996; Rowe & Rundle 2021). As a consequence, sexual selection inflates the genetic variance of fitness components (David *et al*. 2000), eliciting a higher responsiveness to environmental stress in the sex experiencing stronger competition for mates and/or gametes.

Sex-specific stress responses may, however, not only be driven by sex-specific sexual selection but can also evolve as a result of ecological character displacement (De Lisle 2019). According to this model, competition for shared resources generates disruptive selection on resource acquisition traits, which can drive the evolution of ecological sexual dimorphism. Moreover, gamete production is a fundamentally different process in males and females requiring sex-specific physiologies, which in turn may lead to sex-specific responses to changing environments. Among those sex-specific physiological traits, spermatogenesis is often considered to be particularly sensitive to heat stress – an issue that has been extensively studied in mammals including humans (Zorgniotti 1991), and has long been argued to have caused the evolution of external testes (i.e. a scrotum) in several mammalian clades (Moore 1926; Lovegrove 2014). More recent work suggests that heat stress can also have a profound impact on male reproductive performance in ectotherms. For instance, sterilizing temperature of males has been found to be a better predictor of the occurrence of 43 *Drosophila* species than lethal temperature (Parratt *et al*. 2021), suggesting that heat stress on males can impose limits on the geographic distributions of species.

The red flour beetle *Tribolium castaneum*, is a highly tractable study system and an emerging model species for the study of thermal stress responses in ectotherms. In particular, heatwaves of only 5 °C above optimal temperatures have been found to compromise sperm production rate and sperm performance, which translated into higher sensitivity to heat stress in males compared to females under monogamy (Sales *et al*. 2018; Sales, Vasudeva & Gage 2021). In addition, sperm competition trials revealed that heat-stressed males obtain a lower paternity share when competing against control males (Sales *et al*. 2018). However, the role that sexual selection plays as a moderator of sex differences in heat response remains unexplored across ectotherms including *Tribolium* beetles.

In this study we aimed at filling this knowledge gap by testing how sexual selection and thermal stress interact to affect reproductive success of males and females in *T. castaneum*. Specifically, we experimentally manipulated thermal stress by means of simulated heatwaves and permanent heat exposure to contrast heat responses of both sexes in monogamous *versus* polygamous mating system settings. Using this approach, we investigated the following four questions: 1) Does heat stress affect male and female reproductive success differently? 2) Do the sexes recover differently from heat stress? 3) Does the mating system affect heat responses in males and females? 4) Does the mating system interact with sex differences in heat response?

## Material and Methods

### Study system

*Tribolium castaneum* is a highly polygamous species where sexual selection operates especially during postcopulatory episodes in both males and females (Fedina & Lewis 2008). In this experiment we used a highly outbred strain created by TJ in 2020 (called *MTP1*, Montpellier 1). This strain originates from a full-diallel mating design, crossing males and females from five strains (i.e., ‘*Ga1*’, ‘*BRZ5*’, ‘*Japan*’, ‘*Abidjan*’ and ‘*Moliste*’) provided by the Agricultural Research Service of the United States Department of Agriculture (USDA, https://www.ars.usda.gov/). After its creation, the *MTP1* strain has been kept in the lab at a population size of around 1000 individuals with non-overlapping generations.

The *MTP1* strain served as our stock population for wildtype focal individuals and their potential mating partners, whereas beetles from the mutant stock ‘reindeer honey dipper’ (hereafter, *RdHD*) served as competitors during the male and female fitness assays. The *RdHD* strain is characterised by a dominant mutation affecting antenna morphology, which facilitates paternity assessment in the polygamous treatment (i.e., in the setup where *RdHD* individuals acted as competitors of wildtype individuals).

In our lab, located at the Terrain d’expériences of Centre d’Écologie Fonctionnelle et Évolutive in Montpellier, both strains were kept on wheat flour with 5 % dry baker’s yeast at 30 °C in temperature-controlled egg incubators (Ovation 56 Advance; Brinsea Ltd, UK) at a relative humidity of 60%. These were also the conditions of the experiment unless otherwise stated.

### Heat stress treatment

The experiment was carried out between September and December 2020. For logistic reasons, it was run in two consecutive experimental blocks that were one week apart. On day 1 of each block, we transferred approximately 600 wildtype *MTP1* and 400 *RdHD* individuals from the stock cultures to new vials in groups of 50 individuals for egg laying. After three days, we removed all adults and allocated half of the vials from wild-type parents to breed focal individuals whereas the other half was selected to breed potential mating partners. Vials with focal individuals were further randomly assigned to three thermal stress treatments: control (constantly exposed to their adapted temperature of 30 °C), heatwave (exposure to a 5-day heatwave of 40 °C during adulthood prior to mating trials), and permanent heat stress (constantly exposed to 40 °C during development and early adulthood; i.e., from egg laying until mating trials). From day 4 onwards, focal individuals were assigned to the permanent heat exposure (constant exposure to 40 °C). All other individuals (including *RdHD* competitors) remained in control conditions of 30 °C. Between days 21 and 25, we sexed all developed pupae and transferred them to new vials, where they were kept (within their assigned treatment condition) in same-sex groups of 25 individuals so that they remained unmated until the mating trials. On day 39, all individuals had reached adulthood and focal beetles that have been assigned to the heatwave treatment were now exposed to 40 °C for five days.

The thermal optimum of *T. castaneum* is considered to be about 35 °C so that a five-day heat exposure of 40 °C mimics biologically relevant heatwaves for this species (Sales *et al*. 2018). By contrast, permanent exposure to 40 °C is more likely close to the upper critical thermal limit of *T. castaneum* and corresponds to a 5 °C increase of average global surface temperature – a worst-case scenario that is currently projected for 2100 under further increasing emissions of greenhouse gases (IPCC 2021).

Thermal stress treatments were achieved by placing vials with experimental individuals in temperature-controlled egg incubators (Ovation 56 Advance; Brinsea Ltd, UK) at a relative humidity of 60%. In total, we used 6 incubators so that each temperature treatment was replicated in two independent environments.

### Mating trials and fitness assays

From day 44 to 47, after completion of the thermal stress treatments, we conducted mating trials at the control temperature (30 °C), with each trial lasting seven days. Specifically, we applied a full-factorial design and randomly allocated focal males and females from the three thermal stress treatments to two mating system treatments of either monogamous pairs or polygamous groups. Monogamous pairs consisted of two individuals: one focal individual and one wild-type mating partner of the other sex. Polygamous groups comprised ten individuals: one focal individual, five wild-type potential mating partners of the other sex and four *RdHD* competitors of the same sex as the focal. There was hence an equal sex ratio in both treatments. Importantly, there was no scope for competition for mating partners and/or their gametes in monogamous pairs, whereas there was high potential for pre- and postcopulatory sexual selection to operate in polygamous groups. Monogamous and polygamous treatments were kept in 20 mL plastic scintillation vials filled with 5 g of flour medium so that females could lay eggs during the seven-days trial. While we acknowledge that given our experimental design, the mating system treatment was confounded with density (density was five times higher in the polygamous groups), food was provided *ad libitum* to ensure that density effects were negligible.

After the first run of mating trials, we transferred all pairs and groups to fresh vials for a second run of mating trials that also lasted seven days, after which all individuals were discarded. We kept all vials and let all offspring (i.e., eggs laid in the medium) develop until adulthood. Afterwards, all vials were frozen with the adult offspring on day 94 of the experiment. In the following days, reproductive success of focal individuals was assayed by counting the number of adult offspring. In the monogamous treatment, all offspring had wildtype phenotype whereas in polygamous treatment offspring could be either wildtype or *RdHD* phenotype if one of the parents was an *RdHD* competitor (i.e., the mother was *RdHD* competitor in female assays or when sired by *RdHD* competitors in male assays). Fitness of focal individuals from monogamous treatments was quantified as the total number of wildtype offspring. By contrast, fitness of focal individuals from polygamous treatments was quantified as the proportion of wildtype offspring over the total number of offspring (i.e., the sum of wildtype and *RdHD* individuals). Offspring counts from vials obtained in the first mating trials provide fitness estimates immediately after the heat stress treatment (hereafter “fitness assay 1”). By contrast, counts from vials of the second run of mating trials provide fitness estimates after focal individuals had the opportunity to recover from the heat stress treatment for seven days (hereafter “fitness assay 2”). With this setup, we could assess reproductive success of focal individuals over two mating trials (with or without time to recover) to explore sex-specific recovery from thermal stress under contrasting mating systems.

In total, the experiment comprised 175 focal males and 177 focal females (Table S1). On average, each heat stress treatment was replicated 29.3 times for each sex and mating system. However, we obtained a higher sample size from the monogamous than polygamous treatment due to logistical reasons (e.g., more mistakes during the transfer of groups) and because a given replicate was excluded if an individual died during the mating trials (which is five times more likely in groups of ten individuals compared to pairs). Moreover, we observed a higher mortality in heat-stressed individuals, which consequently resulted in higher sample sizes for controls.

### Statistical Analyses

All statistical analyses were carried out in R version 4.0.3 (R Core Team 2021) and included two steps. First, we applied Generalized Linear-Mixed effects Models (GLMMs) implemented in the *lme4* R-package version 1.1.23 (Bates *et al*. 2015) to test for an effect of thermal stress on reproductive success separately for each sex and mating system. For the monogamous treatment, we defined the number of wildtype offspring as the response variable assuming a Poisson error distribution with a log link function. For the polygamous treatment, we defined the number of wildtype offspring and the number of *RdHD* offspring as response variables assuming a binomial error distribution with a logit link function. We fitted heat stress treatment as the only fixed effect together with block, incubator and observation-level as random terms (effects of random terms are shown in Table S2). Observation-level identity was added to account for overdispersion. In addition to tests for an overall treatment effect, we performed *post-hoc* tests to evaluate pair-wise differences between the three heat stress levels.

Second, we computed a selection coefficient to compare the effect of the heat stress treatment between sexes, mating systems, and fitness assays. Specifically, we defined the selection coefficient (*s*) as

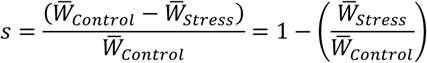

Where 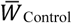 is the mean number of offspring produced under control conditions (i.e., 30 °C) and 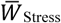 is the mean number of offspring produced under the stressed conditions (i.e., either heatwave of 40 °C for 5 days or permanent exposure to 40 °C). Thus, *s* is the proportional fitness loss due to thermal stress and represents a standardized effect size for the selective advantage of control individuals over heat-stressed individual (Zikovitz & Agrawal 2013; Janicke & Chapuis 2016). We applied bootstrapping with 10,000 iterations using the *boot* R-package version 1.3.28 (Canty & Ripley 2021) to compute 95% confidence intervals for estimates of *s*. Finally, based on the obtained bootstrap samples, we computed the pairwise difference of the selection coefficient Δ*s* and its 95% confidence intervals for the comparison of sexes (Δ*s* _Sex_= *s* _Males_ − *s* _Female_), mating systems (Δ*s* _Mating system_ = *s* _Polygamy_ − *s* _Monogamy_), and fitness assays (Δ*s* _Fitness assay_ = *s* _Fitness assay 1_ − *s* _Fitness assay 2_). We considered estimates of Δ*s* to be significant when confidence intervals did not overlap with zero, whereas estimates centred on zero were considered to provide support for the absence of an effect.

## Results

Heat stress had a substantial impact on reproductive success (Figures 1 and 2; Tables 1 and 2), but the strength of observed effects highly depended on the sex, mating system and fitness assay (Figure 3; Tables 3 and 4). For males, heatwaves impaired male reproductive success when facing competition in the polygamy setting but not under monogamy (Figure 1A and B ; Table 1), which manifested in a significant effect of the mating system (Figure 3; Table 3). The expected negative effect of heatwave on male reproductive success could only be detected immediately after heatwave exposure (fitness assay 1: Figure 1; Table 1), but not after five days of recovery (fitness assay 2: Figure 2; Table 2). In contrast to our findings on the effects of heatwave, permanent heat exposure had a consistent, negative impact across both mating systems (Table 1). Still, similar to the heatwave pattern, the negative effect of permanent heat exposure was stronger under polygamy than under monogamy (Figure 3; Table 4). Importantly, permanent heat stress exposure had a negative effect on males not only immediately after the treatment but also beyond seven days of recovery (Figure 2; Table 2). However, the negative effects were found to be weaker in fitness assay 2 suggesting partial recovery of male reproductive performance (Table 4).

**Table 1.**
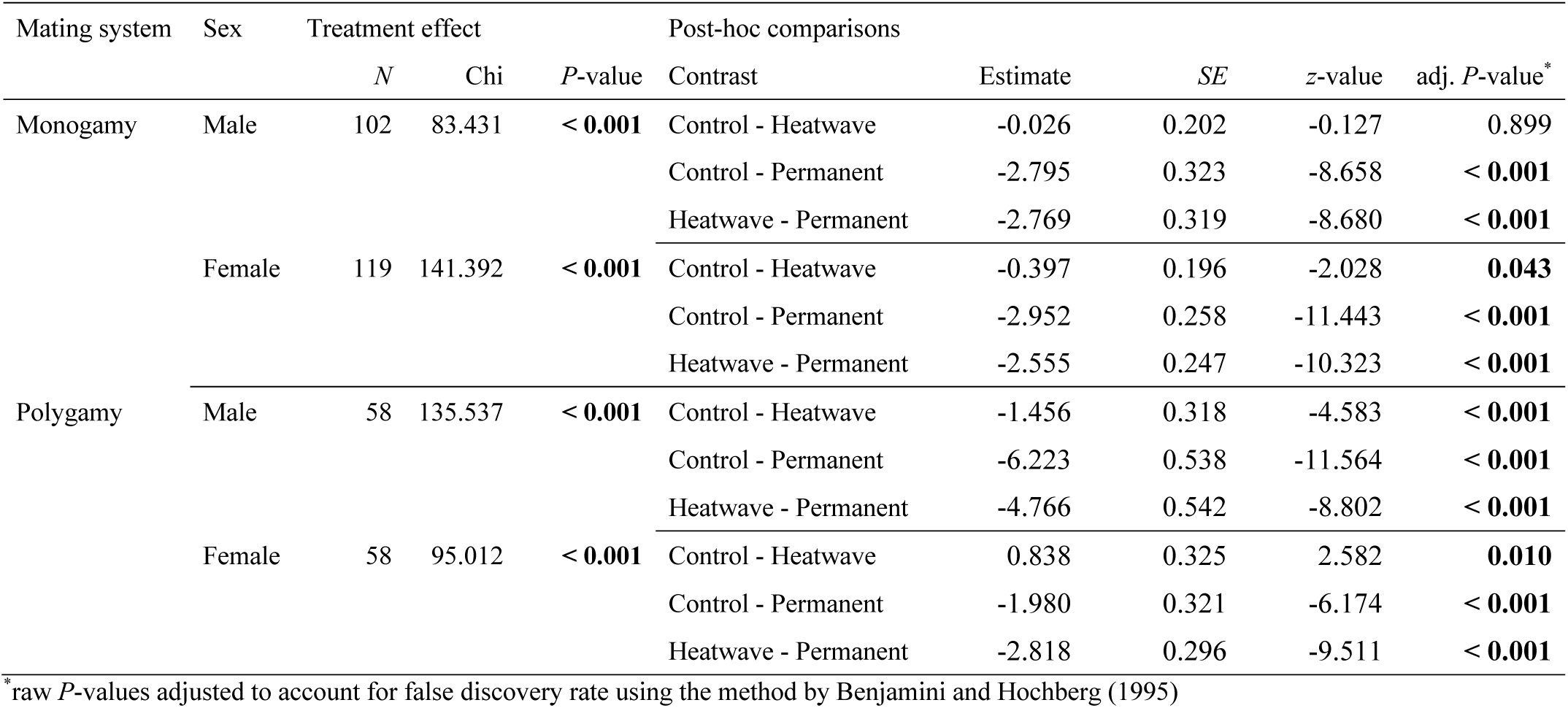
Effect of heat stress on male and female reproductive success immediately after the heat stress treatments (“fitness assay 1”) under monogamy (i.e., one male and one female) and polygamy (i.e., five males and five females). Table shows results obtained from Generalized Linear Mixed-Effects Models testing for an overall treatment effect and all pair-wise comparisons between control (constant 30 °C), heatwave (five days of 40 °C) and permanent heat exposure (constant 40 °C). Statistically significant effects are marked in boldface.

**Table 2.** Effect of heat stress on male and female reproductive success seven days after the heat stress treatments (“fitness assay 2”) under monogamy (i.e., one male and one female) and polygamy (i.e., five males and five females). Table shows results obtained from Generalized Linear Mixed-Effects Models testing for an overall treatment effect and all pair-wise comparisons between control (constant 30 °C), heatwave (five days of 40 °C) and permanent heat exposure (constant 40 °C). Statistically significant effects are marked in boldface.

| Mating System | Sex | Treatment effect |  |  | Post-hoc comparisons |  |  |  |  |
| --- | --- | --- | --- | --- | --- | --- | --- | --- | --- |
| | | *N* | $\chi^2$ | *P*-value | Contrast | Estimate | SE | *z*-value | adj. *P*-value* |
| Monogamy | Male | 102 | 41.140 | < **0.001** | Control - Heatwave | 0.051 | 0.176 | 0.289 | 0.772 |
|  |  |  |  |  | Control - Permanent | -1.460 | 0.250 | -5.845 | < **0.001** |
|  |  |  |  |  | Heatwave - Permanent | -1.511 | 0.247 | -6.129 | < **0.001** |
|  | Female | 119 | 113.552 | < **0.001** | Control - Heatwave | -0.501 | 0.228 | -2.198 | **0.028** |
|  |  |  |  |  | Control - Permanent | -3.099 | 0.299 | -10.354 | < **0.001** |
|  |  |  |  |  | Heatwave - Permanent | -2.598 | 0.287 | -9.037 | < **0.001** |
| Polygamy | Male | 58 | 96.953 | < **0.001** | Control - Heatwave | -0.270 | 0.399 | -0.676 | 0.499 |
|  |  |  |  |  | Control - Permanent | -4.388 | 0.476 | -9.225 | < **0.001** |
|  |  |  |  |  | Heatwave - Permanent | -4.118 | 0.478 | -8.609 | < **0.001** |
|  | Female | 58 | 52.996 | < **0.001** | Control - Heatwave | 0.952 | 0.426 | 2.234 | **0.025** |
|  |  |  |  |  | Control - Permanent | -1.563 | 0.428 | -3.652 | < **0.001** |
|  |  |  |  |  | Heatwave - Permanent | -2.515 | 0.347 | -7.242 | < **0.001** |

\*raw *P*-values adjusted to account for false discovery rate using the method by Benjamini and Hochberg (1995)

**Table 3.** Effect of heatwave treatment on male and female reproductive success. Table shows bootstrapped selection coefficients together with 95% intervals (in brackets) estimating the selective advantage of control individuals (constant 30 °C) over heat stressed individuals (five days of heatwave of 40 °C). Bootstrapped differences Δ*s* between mating systems, sexes and fitness assays are marked in boldface whenever confidence intervals do not include zero.

| Fitness assay | Mating system | Male | | Female | | $\Delta s_{\text{Sex}}$ | |
| --- | --- | --- | --- | --- | --- | --- | --- |
| Fitness assay 1 | Monogamy | 0.060 | (-0.175, 0.249) | 0.232 | (0.072, 0.371) | -0.171 | (-0.445, 0.077) |
|  | Polygamy | 0.576 | (0.420, 0.702) | -0.715 | (-1.323, -0.248) | **1.290** | **(0.801, 1.917)** |
| | $\Delta s_{\text{Mating system}}$ | **0.515** | **(0.267, 0.781)** | **-0.947** | **(-1.572, -0.452)** | | |
| Fitness assay 2 | Monogamy | 0.015 | (-0.179, 0.181) | 0.308 | (0.151, 0.449) | **-0.294** | **(-0.528, -0.061)** |
|  | Polygamy | 0.104 | (-0.103, 0.295) | -0.829 | (-1.617, -0.288) | **0.934** | **(0.345, 1.743)** |
| | $\Delta s_{\text{Mating system}}$ | 0.090 | (-0.175, 0.358) | **-1.138** | **(-1.945, -0.572)** | | |
| $\Delta s_{\text{Fitness assay}}$ | Monogamy | 0.046 | (-0.238, 0.317) | -0.077 | (-0.285, 0.138) | | |
|  | Polygamy | **0.471** | **(0.226, 0.714)** | 0.115 | (-0.703, 1.045) |  |  |

**Table 4.** Effect of permanent heat exposure on male and female reproductive success. Table shows bootstrapped selection coefficients *s* together with 95% intervals (in brackets) estimating the selective advantage of control individuals (constant 30 °C) over heat-stressed individuals (permanent heat exposure to 40 °C). Bootstrapped differences Δ*s* between mating systems, sexes and fitness assays are marked in boldface whenever confidence intervals do not include zero.

| Fitness assay | Mating system | Male | | Female | | $\Delta s_{\text{Sex}}$ | |
| --- | --- | --- | --- | --- | --- | --- | --- |
| Fitness assay 1 | Monogamy | 0.850 | (0.716, 0.960) | 0.886 | (0.806, 0.948) | -0.036 | (-0.185, 0.101) |
|  | Polygamy | 0.991 | (0.976, 1.000) | 0.798 | (0.677, 0.893) | **0.193** | **(0.098, 0.314)** |
| | $\Delta s_{\text{Mating system}}$ | **0.141** | **(0.031, 0.276)** | -0.088 | (-0.224, 0.035) | | |
| Fitness assay 2 | Monogamy | 0.514 | (0.214, 0.773) | 0.873 | (0.779, 0.947) | **-0.359** | **(-0.671, -0.086)** |
|  | Polygamy | 0.891 | (0.782, 0.976) | 0.661 | (0.426, 0.837) | **0.230** | **(0.023, 0.484)** |
| | $\Delta s_{\text{Mating system}}$ | **0.377** | **(0.100, 0.689)** | **-0.212** | **(-0.457, -0.013)** | | |
| $\Delta s_{\text{Fitness assay}}$ | Monogamy | **0.336** | **(0.047, 0.659)** | 0.013 | (-0.096, 0.126) | | |
|  | Polygamy | **0.100** | **(0.015, 0.209)** | 0.137 | (-0.077, 0.396) |  |  |

**Figure 1.**
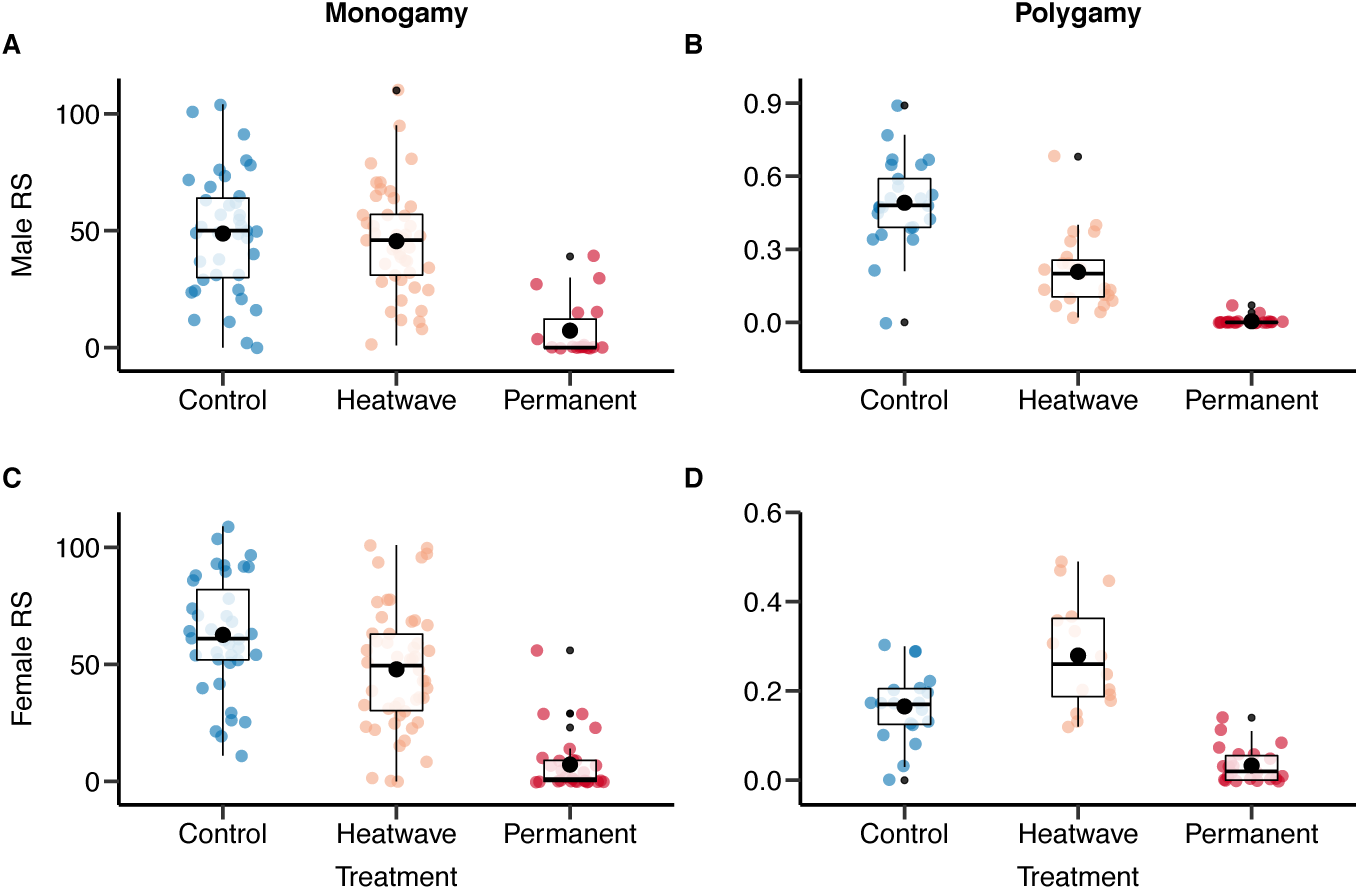
Comparison of reproductive success between heat stress treatments when measured immediately after heat exposure (“fitness assay 1”). Boxplots are shown for males (A, B) and females (C, D) under monogamy (A, C) and polygamy (B, D). Heat stress treatment includes control (constant 30 °C), heatwave (five days of 40 °C), and permanent heat stress exposure (constant 40 °C). Boxplots show the 25^th^ percentile, the median, and the 75^th^ percentile and whiskers denote the 5^th^ and the 95^th^ percentiles. Filled black circles represent the means.

**Figure 2.**
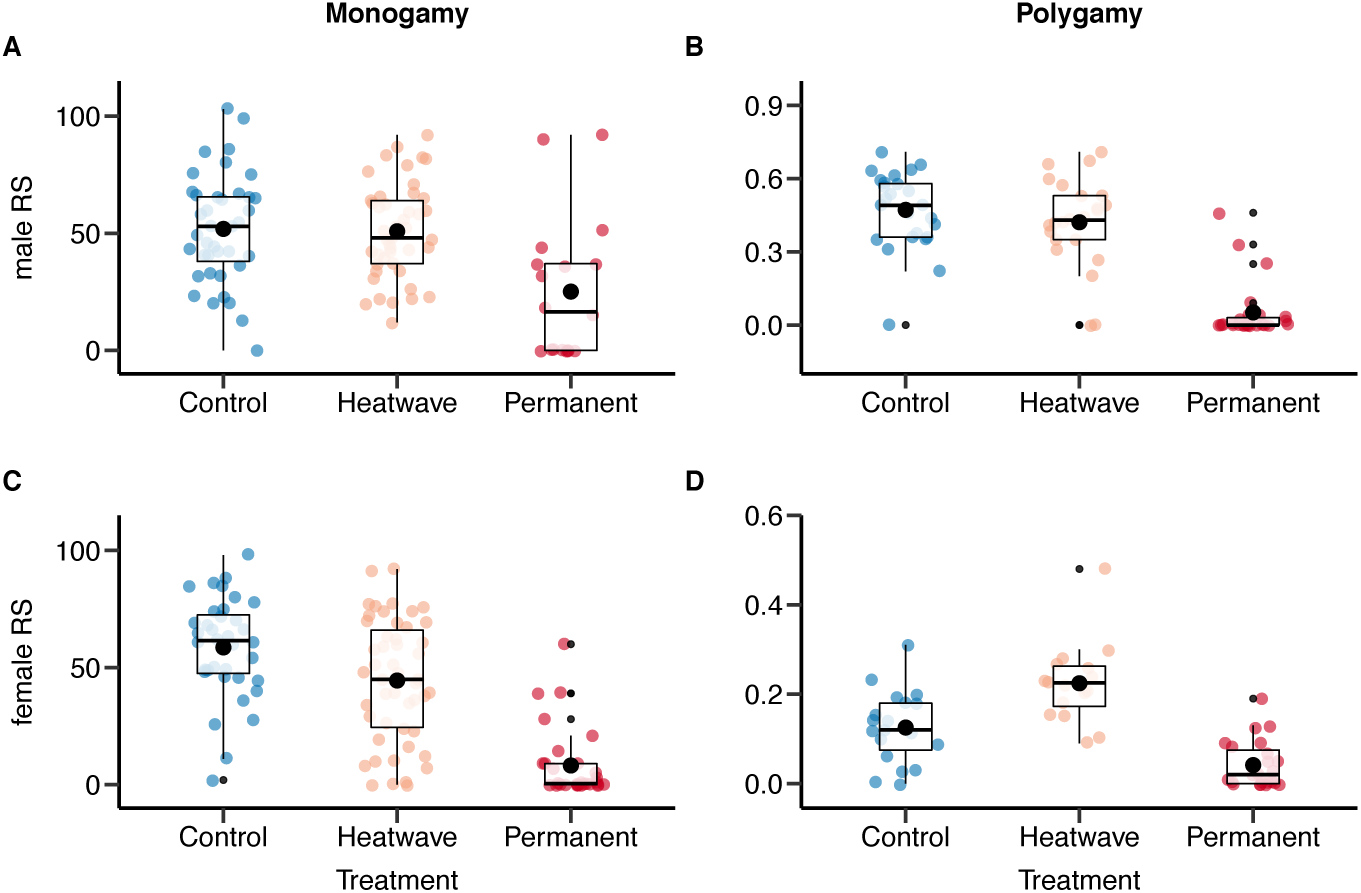
Comparison of reproductive success between heat stress treatments when measured seven days after heat exposure (“fitness assay 2”). Boxplots are shown for males (A, B) and females (C, D) under monogamy (A, C) and polygamy (B, D). Heat stress treatment includes control (constant 30 °C), heatwave (five days of 40 °C), and permanent heat stress exposure (constant 40 °C). Boxplots show the 25^th^ percentile, the median, and the 75^th^ percentile and whiskers denote the 5^th^ and the 95^th^ percentiles. Filled black circles represent the means.

**Figure 3.**
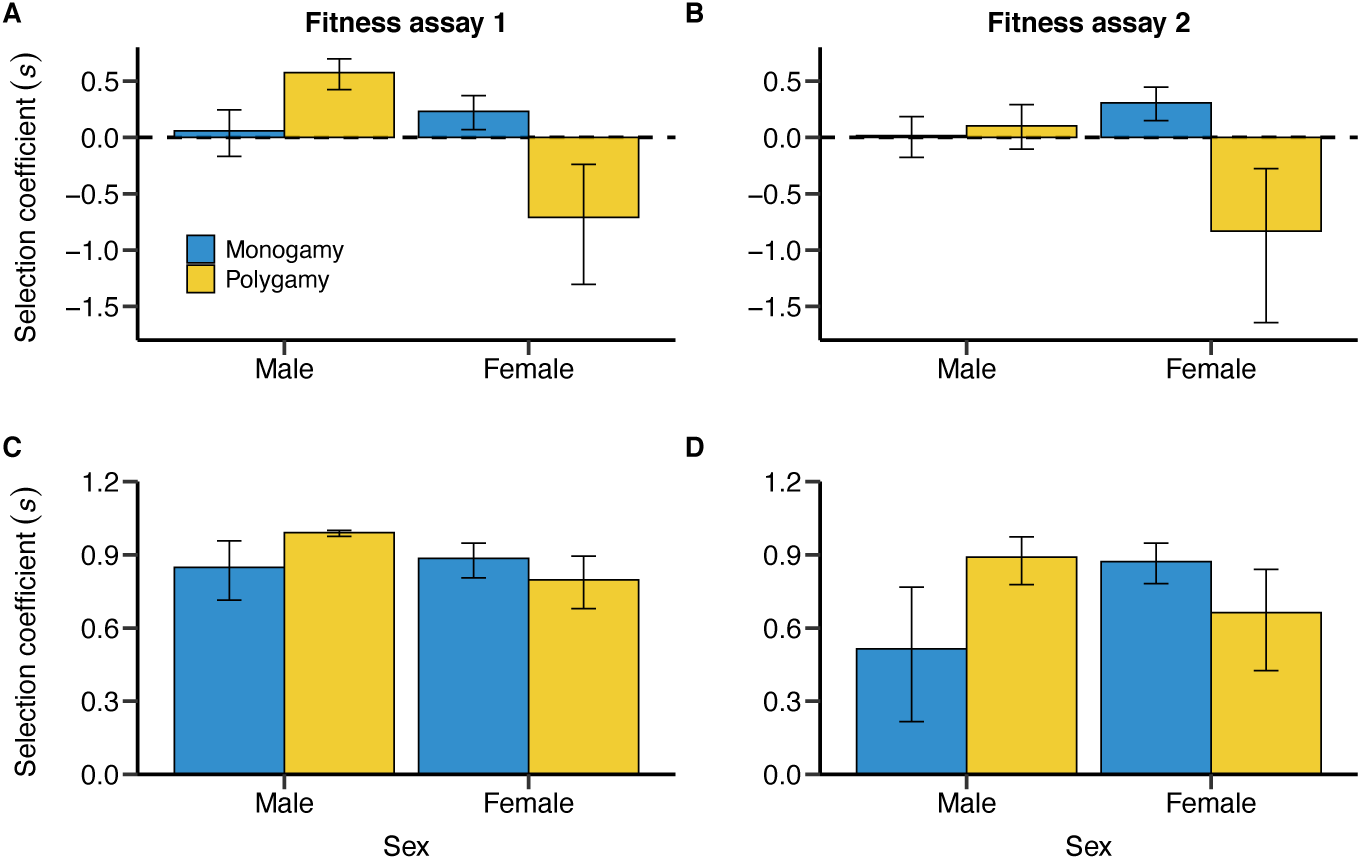
Effects of mating system on selection coefficients (*s*) estimating the selective advantage of control individuals over heat stressed individuals. Selection coefficients are shown for males and females under monogamy (blue) and polygamy (yellow). Comparisons shown for heatwave (A, B) and permanent heat exposure (C, D) when measured directly after the treatment (A, C; “fitness assay 1”) and seven days after the treatment (B, D; “fitness assay 2”). Error bars denote bootstrapped means together with 95% confidence intervals.

In females, heatwave had a weak but significant negative effect on reproductive success under monogamy (Figure 1C; Table 1). In the polygamy treatment, however, we observed a higher reproductive output of females exposed to the heatwave (Figure 1D; Table 1). Thus, the mating system had a strong effect on heatwave response in females, changing its direction (Figure 3; Table 3). By contrast, permanent heat exposure had a negative effect under both mating systems (Figure 2C and D). This negative effect was generally stronger in the monogamy treatment but was found to be significantly different between mating systems only in fitness assay 2 (Table 4). Overall, we found that the observed effects of heatwave and permanent heat exposure on females remained largely constant over the two fitness assays (Tables 3 and 4).

Finally, heat stress also had sex-specific effects. Under monogamy, both heatwave and permanent heat exposure had a more negative effect on females than on males, but this sex difference was only found to be statistically significant in fitness assay 2 (Tables 3 and 4). By contrast, under polygamy, both heat stress treatments had a stronger negative effect on males than on females (Tables 3 and 4).

## Discussion

Our study provides compelling evidence for the role of sexual selection as a moderator of sex-specific heat stress responses in the form of four main findings. First, heat stress affected reproductive performance of males and females. Second, mating systems altered heat-stress responses in both sexes. Remarkably, the observed effect of sexual selection on heatwave responses was found to have opposing directions for males and females. Third, males showed full recovery from heatwave and partial recovery from permanent heat stress exposure. By contrast beneficial and detrimental effects on females had long-term effects with no signs of deterioration or recovery, respectively. And fourth, the sex difference in heat response was highly dependent on the mating system. While heat stress had a stronger effect on females in the monogamy setting, we observed a stronger sensitivity to heat stress in males under polygamy.

### Effect of heat stress on male and female reproductive success

For males, we detected a non-significant, negative effect of heatwave in the monogamy treatment but a reduction of 57.6% in reproductive success under polygamy, i.e., under conditions allowing for pre- and postcopulatory sexual selection to operate. Therefore, the presence of sexual selection is associated with detrimental effects of heat stress on males. Permanent heat exposure impaired male reproductive success drastically in both mating systems, but similar to our findings on heatwaves, the effect was magnified in the polygamy treatment in which we observed an almost complete reproductive failure (i.e., a reduction of 99.1%). Overall, these findings are consistent with results of a previous study demonstrating a negative effect of a five-days heatwave on male sperm competitiveness inferred from assays in which focal males were paired with an already mated female (Sales *et al*. 2018). Interestingly, Sales *et al*. (2018) also investigated the underlying mechanisms causing the observed heat response, and showed that heatwaves act by lowering mating rates and reducing both sperm numbers and sperm viability.

Contrary to our results, Sales *et al*. (2018) and Sales, Vasudeva and Gage (2021) observed a reduction of male reproductive success of 21.3% after heatwaves of 40 °C under monogamy. We detected only a marginal and not statistically significant tendency for a reduction of male reproductive success of 6.1%. We suspect that two major differences in the experimental design between our and the previous studies may have contributed to the contrasting findings under the monogamy setting. First, both studies used different stock cultures, which may have differed in the standing genetic variation for heat resistance. However, both studies used highly outbred strains, which renders this hypothesis unlikely to be true. Second, the two previous studies applied an experimental setup in which mating trials took place one day after the heatwave treatment, lasted only two days and were temporally separated from the fitness assays (Sales *et al*. 2018; Sales, Vasudeva & Gage 2021). By contrast, in our experiment, mating trials and fitness assays were coupled and the first fitness assay started immediately after the thermal stress treatment lasting for seven days. Consequently, in our setup, male reproductive success was recorded directly after heatwave exposure but males could recover longer from heat stress during the mating trials compared to the setup used by Sales *et al*. (2018), which might have mitigated the detrimental effects reported by earlier work.

For females, we observed a lowered reproductive success of individuals experiencing heatwave (23.2 % reduction) or permanent thermal stress (88.6 % reduction) under monogamy. These findings also contradict results of previous studies that did not detect significant differences between females exposed to 30 °C and 40 °C arguing that *T. castaneum* has its thermal optimum at 35 °C (Sales *et al*. 2018; Sales, Vasudeva & Gage 2021). Nevertheless, evidence from other studies on *T. castaneum* are in line with our findings, documenting a lowered female fecundity under monogamy at 35 °C or 37 °C compared to 30 °C or 33 °C, respectively (Koch & Guillaume 2020; Fischer *et al*. 2021). Once again, differences in findings may result from slight differences in the experimental designs. Regarding the results on heatwave exposure, we suspect that detrimental effects on female fecundity may only play out when oviposition takes place directly after the heatwave and may be smaller when fitness assays start three days after the heatwave treatment (Sales *et al*. 2018).

Presumably the most surprising finding of our study is the observed positive net effect of heatwave on female reproductive success under polygamy. Even though permanent heat exposure had a negative effect on female reproductive success under polygamy, its magnitude was lower compared to the negative effect detected under monogamy. Thus, our findings suggest that females may benefit from multiple mating under elevated temperatures. Experimental studies testing for fitness consequences of polyandry in *T. castaneum* provided mixed results, but overall suggest that females obtain indirect rather than direct benefits (Bernasconi & Keller 2001; Pai & Yan 2002; Pai, Bennett & Yan 2005; Pai, Feil & Yan 2007; Pai & Yan 2020; Matsumura, Miyatake & Yasui 2021). As such, the question of how potential benefits of polyandry depend on the thermal conditions remains largely unexplored. Interestingly, Grazer and Martin (2012) exposed female *T. castaneum* to ambient temperatures of 30 °C *versus* 34 °C and observed an increase in reproductive success at elevated temperatures under polygamy but not under monogamy – similar to our findings. They argued that seminal fluids may mitigate detrimental effects of temperature and desiccation stress (Poiani 2006) (Poiani 2006; Edvardsson 2007), because they contain important substances such as nutrients and water, that are a limiting factor for female reproductive success, particularly under elevated temperatures (Grazer & Martin 2012). Red flour beetles only acquire water indirectly via air humidity and food, but not directly by drinking. Hence, polygamous females may benefit from multiple mating under elevated temperatures by receiving water- and nutrient-rich spermatophores as documented in other insects such as bruchid beetles (Edvardsson 2007) and bush-crickets (Gwynne & Simmons 1990). Further work will be required to identify the underlying processes explaining the observed interaction between thermal stress and mating system. Moreover, our results illustrate the need to study heat response in females not only under monogamy but also in settings allowing for sexual selection to operate. Sexual selection is widespread not only in males but also in females (Hare & Simmons 2019; Fromonteil *et al*. 2021), but its effect on female stress responses is still poorly understood.

Finally, our study also highlights an important aspect regarding the temporal persistence of the observed effects. Specifically, we found evidence that males have the capacity to recover from heat stress (fully or partly depending on the duration of the heat stress treatment), whereas effects on females remained constant along the tested time window of two weeks after heat exposure. These findings largely coincide with findings by Sales, Vasudeva and Gage (2021) who detected full recovery of male reproductive performance 25 days after heat exposure.

### Effect of sexual selection on sex-specific heat response

Our study demonstrates that polygamy triggers a negative heat response in males but relaxes detrimental effects of heat stress in females. This striking finding suggests that sexual selection can reverse the sex differences in heat sensitivity. Thermal stress has often been argued to have stronger effects on males compared to females but experimental studies in arthropods testing for sex-specific heat responses provided mixed results ranging from a male-bias (Sales *et al*. 2018; Walsh *et al*. 2021) to no sex differences (Piyaphongkul, Pritchard & Bale 2012) and female-biased thermal sensitivity (Janowitz & Fischer 2011). Our results suggest that mating system might be an important determinant of intra- and inter-specific variation of sex differences in heat sensitivity. In a recent study, Baur *et al*. (2022) tested for a short-term evolutionary response in heat sensitivity under varying mating systems in the seed beetle *Callosobruchus maculatus*. Their findings suggest that populations evolving under polygamy show a higher sensitivity to heat stress in females compared to males when reproductive success was assayed under monogamy. Thus, our findings together with the results from Baur *et al*. (2022) indicate that mating system can be a potent driver for sex differences in heat response at the phenotypically plastic and micro-evolutionary scale.

Sex differences in heat response are important to project the adaptive responses of populations to increased temperatures, which is a pressing question in the face of current global warming and increased frequency of extreme climatic events. Theory predicts that sexual selection can accelerate adaptation to changing environments when two conditions are fulfilled. First, sexual selection and natural selection need to be aligned meaning that sexual selection favours genetic variants that are also beneficial under natural selection (Rowe & Rundle 2021). Second, net selection must be stronger on males compared to females so that populations can purge deleterious alleles at a low demographic cost (Whitlock & Agrawal 2009; Winkler *et al*. 2021). In *T. castaneum*, there is compelling evidence for an alignment of natural and sexual selection (Lumley *et al*. 2015; Godwin *et al*. 2020), however it remains to be tested whether alleles that are beneficial under thermal stress in males are also advantageous under thermal stress in females. Our study suggests that sexual selection (i.e., polygamy) intensifies net selection on heat-resistance in males but not in females. Hence, adaptation to increased temperatures may indeed be mediated by sexual selection on males as documented for other insects (Plesnar-Bielak *et al*. 2012; Parrett & Knell 2018).

### Conclusions

Our findings demonstrate that the intensity of sexual selection plays a pivotal role in predicting heat responses in both sexes. We found that polygamy magnifies detrimental effects of heat stress in males but relaxes negative effects on females. Thus, sexual selection may reverse the sex difference in thermal sensitivity and thereby the sex-specific strength of selection on alleles associated with heat resistance. A better understanding of how populations adapt to rising temperatures and increased frequency of extreme climatic events will critically depend on more detailed knowledge on the thermal optima under varying levels of sexual selection. Importantly, experimental studies testing for an effect of sexual selection on heat responses should not only focus on males but also include females. Finally, quantitative genetics and genomic approaches have the potential to examine whether heat stress favours the same genetic variants in male and females, which is essential to evaluate the role of sexual selection for modulating adaptation to climate change.

## Acknowledgements

We are grateful to Thierry Mathieu and Pierrick Aury from the LabEx CeMEB platform “Terrain d’Expériences” in Montpellier for logistic support.

## References

Angilletta, M.J. (2009) Thermal adaptation: a theoretical and empirical synthesis. Oxford University Press, Inc., New York.

Bates, D., Maechler, M., Bolker, B. & Walker, S. (2015) Fitting Linear Mixed-Effects Models using lme4. Journal of Statistical Software, 67, 1–48.

Baur, J., Jagusch, D., Michalak, P., Koppik, M. & Berger, D. (2022) The mating system affects the temperature sensitivity of male and female fertility. Functional Ecology, 36, 92–106.

Bellard, C., Bertelsmeier, C., Leadley, P., Thuiller, W. & Courchamp, F. (2012) Impacts of climate change on the future of biodiversity. Ecology Letters, 15, 365–377.

Benjamini, Y. & Hochberg, Y. (1995) Controlling the false discovery rate - a practical and powerful approach to multiple testing. Journal of the Royal Statistical Society Series B-Statistical Methodology, 57, 289–300.

Bernasconi, G. & Keller, L. (2001) Female polyandry affects their sons’ reproductive success in the red flour beetle Tribolium castaneum. Journal of Evolutionary Biology, 14, 186–193.

Bonduriansky, R., Maklakov, A., Zajitschek, F. & Brooks, R. (2008) Sexual selection, sexual conflict and the evolution of ageing and life span. Functional Ecology, 22, 443–453.

Canty, A. & Ripley, B.D. (2021) boot: Bootstrap R (S-Plus) functions. R package version 1.3-28.

Coumou, D. & Rahmstorf, S. (2012) A decade of weather extremes. Nature Climate Change, 2, 491–496.

David, P., Bjorksten, T., Fowler, K. & Pomiankowski, A. (2000) Condition-dependent signalling of genetic variation in stalk-eyed flies. Nature, 406, 186–188.

De Lisle, S.P. (2019) Understanding the evolution of ecological sex differences: Integrating character displacement and the Darwin-Bateman paradigm. Evolution Letters, 3, 434–447.

Edvardsson, M. (2007) Female Callosobruchus maculatus mate when they are thirsty: resource-rich ejaculates as mating effort in a beetle. Animal Behaviour, 74, 183–188.

Fedina, T.Y. & Lewis, S.M. (2008) An integrative view of sexual selection in Tribolium flour beetles. Biological Reviews, 83, 151–171.

Fischer, E.M. & Knutti, R. (2015) Anthropogenic contribution to global occurrence of heavy-precipitation and high-temperature extremes. Nature Climate Change, 5, 560–564.

Fischer, K., Kreyling, J., Beaulieu, M., Beil, I., Bog, M., Bonte, D., Holm, S., Knoblauch, S., Koch, D., Muffler, L., Mouginot, P., Paulinich, M., Scheepens, J.F., Schiemann, R., Schmeddes, J., Schnittler, M., Uhl, G., van der Maaten-Theunissen, M., Weier, J.M., Wilmking, M., Weigel, R. & Gienapp, P. (2021) Species-specific effects of thermal stress on the expression of genetic variation across a diverse group of plant and animal taxa under experimental conditions. Heredity, 126, 23–37.

Fromonteil, S., Winkler, L., Marie-Orleach, L. & Janicke, T. (2021) Sexual selection in females across the animal tree of life. bioRxiv, doi: https://doi.org/10.1101/2021.1105.1125.445581

Garcia-Roa, R., Garcia-Gonzalez, F., Noble, D.W.A. & Carazo, P. (2020) Temperature as a modulator of sexual selection. Biological Reviews, 95, 1607–1629.

Godwin, J.L., Lumley, A.J., Michalczyk, Ł., Martin, O.Y. & Gage, M.J. (2020) Mating patterns influence vulnerability to the extinction vortex. Global Change Biology, 26, 4226–4239.

Grazer, V.M. & Martin, O.Y. (2012) Elevated temperature changes female costs and benefits of reproduction. Evolutionary Ecology, 26, 625–637.

Gwynne, D.T. & Simmons, L.W. (1990) Experimental reversal of courtship roles in an insect. Nature, 346, 172–174.

Hämäläinen, A., Immonen, E., Tarka, M. & Schuett, W. (2018) Evolution of sex-specific pace-of-life syndromes: causes and consequences. Behavioral Ecology and Sociobiology, 72, 1–15.

Hare, R.M. & Simmons, L.W. (2019) Sexual selection and its evolutionary consequences in female animals. Biological Reviews, 94, 929–956.

Huey, R.B., Kearney, M.R., Krockenberger, A., Holtum, J.A.M., Jess, M. & Williams, S.E. (2012) Predicting organismal vulnerability to climate warming: roles of behaviour, physiology and adaptation. Philosophical Transactions of the Royal Society B-Biological Sciences, 367, 1665–1679.

Huey, R.B. & Kingsolver, J.G. (1993) Evolution of resistance to high temperature in ectotherms. The American Naturalist, 142, S21–S46.

IPCC (2021) Climate Change 2021. The Physical Science Basis. Contribution of Working Group I to the Sixth Assessment Report of the Intergovernmental Panel on Climate Change. Cambridge University Press.

Janicke, T. & Chapuis, E. (2016) Condition-dependence of male and female reproductive success: insights from a hermaphrodite. Ecology and evolution, 6, 830–841.

Janicke, T., Häderer, I.K., Lajeunesse, M.J. & Anthes, N. (2016) Darwinian sex roles confirmed across the animal kingdom. Science Advances, 2, e1500983.

Janowitz, S.A. & Fischer, K. (2011) Opposing effects of heat stress on male versus female reproductive success in Bicyclus anynana butterflies. Journal of Thermal Biology, 36, 283–287.

Jones, A.G. & Ratterman, N.L. (2009) Mate choice and sexual selection: what have we learned since Darwin? Proceedings of the National Academy of Sciences of the United States of America, 106, 10001–10008.

Koch, E.L. & Guillaume, F. (2020) Additive and mostly adaptive plastic responses of gene expression to multiple stress in Tribolium castaneum. PLoS genetics, 16, e1008768.

Lovegrove, B. (2014) Cool sperm: why some placental mammals have a scrotum. Journal of Evolutionary Biology, 27, 801–814.

Lumley, A.J., Michalczyk, L., Kitson, J.J.N., Spurgin, L.G., Morrison, C.A., Godwin, J.L., Dickinson, M.E., Martin, O.Y., Emerson, B.C., Chapman, T. & Gage, M.J.G. (2015) Sexual selection protects against extinction. Nature, 522, 470–473.

Matsumura, K., Miyatake, T. & Yasui, Y. (2021) An empirical test of the bet-hedging polyandry hypothesis: Female red flour beetles avoid extinction via multiple mating. Ecology and evolution, 11, 5295–5304.

Moore, C.R. (1926) The biology of the mammalian testis and scrotum. Quarterly Review of Biology, 1, 4–50.

Pai, A., Bennett, L. & Yan, G.Y. (2005) Female multiple mating for fertility assurance in red flour beetles (Tribolium castaneum). Canadian Journal of Zoology, 83, 913–919.

Pai, A., Feil, S. & Yan, G. (2007) Variation in polyandry and its fitness consequences among populations of the red flour beetle, Tribolium castaneum. Evolutionary Ecology, 21, 687–702.

Pai, A.D. & Yan, G.Y. (2020) Long-term study of female multiple mating indicates direct benefits in Tribolium castaneum. Entomologia Experimentalis Et Applicata, 168, 398–406.

Pai, A.T. & Yan, G.Y. (2002) Polyandry produces sexy sons at the cost of daughters in red flour beetles. Proceedings of the Royal Society B-Biological Sciences, 269, 361–368.

Parmesan, C. (2006) Ecological and evolutionary responses to recent climate change. Annual Review of Ecology Evolution and Systematics, 37, 637–669.

Parratt, S.R., Walsh, B.S., Metelmann, S., White, N., Manser, A., Bretman, A.J., Hoffmann, A.A., Snook, R.R. & Price, T.A. (2021) Temperatures that sterilize males better match global species distributions than lethal temperatures. Nature Climate Change, 11, 481–484.

Parrett, J.M. & Knell, R.J. (2018) The effect of sexual selection on adaptation and extinction under increasing temperatures. Proceedings of the Royal Society B-Biological Sciences, 285, 7.

Pilakouta, N. & Ålund, M. (2021) Sexual selection and environmental change: what do we know and what comes next? Current Zoology, 67, 293–298.

Piyaphongkul, J., Pritchard, J. & Bale, J. (2012) Can tropical insects stand the heat? A case study with the brown planthopper Nilaparvata lugens (Stål). PloS ONE, 7, e29409.

Plesnar-Bielak, A., Skrzynecka, A.M., Prokop, Z.M. & Radwan, J. (2012) Mating system affects population performance and extinction risk under environmental challenge. Proceedings of the Royal Society B: Biological Sciences, 279, 4661–4667.

Poiani, A. (2006) Complexity of seminal fluid: a review. Behavioral Ecology and Sociobiology, 60, 289–310.

R Core Team (2021) A language and environment for statistical computing. R Foundation for Statistical Computing, Vienna, Austria. URL http://www.R-project.org/.

Rowe, L. & Houle, D. (1996) The lek paradox and the capture of genetic variance by condition dependent traits. Proceedings of the Royal Society of London Series B-Biological Sciences, 263, 1415–1421.

Rowe, L. & Rundle, H.D. (2021) The alignment of natural and sexual selection. Annual Review of Ecology, Evolution, and Systematics, 52.

Sales, K., Vasudeva, R., Dickinson, M.E., Godwin, J.L., Lumley, A.J., Michalczyk, L., Hebberecht, L., Thomas, P., Franco, A. & Gage, M.J.G. (2018) Experimental heatwaves compromise sperm function and cause transgenerational damage in a model insect. Nature Communications, 9, 11.

Sales, K., Vasudeva, R. & Gage, M.J. (2021) Fertility and mortality impacts of thermal stress from experimental heatwaves on different life stages and their recovery in a model insect. Royal Society open science, 8, 201717.

Schärer, L., Rowe, L. & Arnqvist, G. (2012) Anisogamy, chance and the evolution of sex roles. Trends in Ecology & Evolution, 27, 260–264.

Stoehr, A.M. & Kokko, H. (2006) Sexual dimorphism in immunocompetence: what does life-history theory predict? Behavioral Ecology, 17, 751–756.

Tarka, M., Guenther, A., Niemelä, P.T., Nakagawa, S. & Noble, D.W. (2018) Sex differences in life history, behavior, and physiology along a slow-fast continuum: a meta-analysis. Behavioral Ecology and Sociobiology, 72, 1–13.

Tattersall, G.J., Sinclair, B.J., Withers, P.C., Fields, P.A., Seebacher, F., Cooper, C.E. & Maloney, S.K. (2012) Coping with thermal challenges: physiological adaptations to environmental temperatures. Comprehensive Physiology, 2, 2151–2202.

Ummenhofer, C.C. & Meehl, G.A. (2017) Extreme weather and climate events with ecological relevance: a review. Philosophical Transactions of the Royal Society B-Biological Sciences, 372.

Walsh, B.S., Mannion, N.L., Price, T.A. & Parratt, S.R. (2021) Sex-specific sterility caused by extreme temperatures is likely to create cryptic changes to the operational sex ratio in Drosophila virilis. Current Zoology, 67, 341–343.

Walsh, B.S., Parratt, S.R., Hoffmann, A.A., Atkinson, D., Snook, R.R., Bretman, A. & Price, T.A. (2019) The impact of climate change on fertility. Trends in Ecology & Evolution, 34, 249–259.

Whitlock, M.C. & Agrawal, A.F. (2009) Purging the genome with sexual selection: reducing mutation load through selection on males. Evolution, 63, 569–582.

Winkler, L., Moiron, M., Morrow, E.H. & Janicke, T. (2021) Stronger net selection on males across animals. Elife, 10, e68316.

Zikovitz, A.E. & Agrawal, A.F. (2013) The condition dependency of fitness in males and females: the fitness consequences of juvenile diet assessed in environments differing in key adult resources. Evolution, 67, 2849–2860.

Zorgniotti, A.W. (1991) Temperature and environmental effects on the testis. Springer.

