## Supplementary Information for "Sexual selection moderates heat stress response in males and females"

This Supplementary Information file includes:

Tables S1 and S2

**Table S1. Overview of sample sizes.** Number of focal male and female individuals (*N*) in each mating system and heat stress treatments.

| Sex | Mating System | Temperature | *N* |
|---|---|---|---|
| Male | Monogamy | Control | 39 |
| | | Heatwave | 45 |
| | | Permanent | 18 |
| | Polygamy | Control | 25 |
| | | Heatwave | 23 |
| | | Permanent | 25 |
| Female | Monogamy | Control | 39 |
| | | Heatwave | 50 |
| | | Permanent | 30 |
| | Polygamy | Control | 19 |
| | | Heatwave | 16 |
| | | Permanent | 23 |

**Table S2. Effects of random terms (Block and Incubator) on male and female reproductive success shown for both fitness assays (1 and 2) and mating systems (Monogamy and Polygamy).** Table shows results obtained from Generalized Linear Mixed-Effects Models testing for an overall treatment effect (see Main Text for fixed effects). Statistically significant effects are marked in boldface.

| Fitness assay | Mating system | Sex | Term | *df* | $\chi^2$ | *P*-value |
|---|---|---|---|---|---|---|
| 1 | Monogamy | Male | Block | 1 | < 0.001 | > 0.999 |
| | | | Incubator | 1 | < 0.001 | > 0.999 |
| | | Female | Block | 1 | < 0.001 | > 0.999 |
| | | | Incubator | 1 | < 0.001 | > 0.999 |
| | Polygamy | Male | Block | 1 | < 0.001 | > 0.999 |
| | | | Incubator | 1 | < 0.001 | > 0.999 |
| | | Female | Block | 1 | 1.099 | 0.295 |
| | | | Incubator | 1 | 0.365 | 0.546 |
| 2 | Monogamy | Male | Block | 1 | 2.348 | 0.125 |
| | | | Incubator | 1 | < 0.001 | > 0.999 |
| | | Female | Block | 1 | 5.827 | 0.016 |
| | | | Incubator | 1 | < 0.001 | > 0.999 |
| | Polygamy | Male | Block | 1 | 0.124 | 0.724 |
| | | | Incubator | 1 | < 0.001 | > 0.999 |
| | | Female | Block | 1 | < 0.001 | > 0.999 |
| | | | Incubator | 1 | 1.039 | 0.308 |
